# A functionally divergent intrinsically disordered region underlying the conservation of stochastic signaling

**DOI:** 10.1101/2021.06.01.446530

**Authors:** Ian S Hsu, Bob Strome, Emma Lash, Nicole Robbins, Leah E Cowen, Alan M Moses

**Affiliations:** Department of Cell & Systems Biology, University of Toronto; Department of Molecular Genetics, University of Toronto; Department of Computer Science, University of Toronto

## Abstract

Stochastic signaling dynamics expand living cells’ information processing capabilities. An increasing number of studies report that regulators encode information in their pulsatile dynamics. The evolutionary mechanisms that lead to complex signaling dynamics remain uncharacterized, perhaps because key interactions of signaling proteins are encoded in intrinsically disordered regions (IDRs), whose evolution is difficult to analyze. Here we focused on the stochastic pulsing dynamics of Crz1, a transcription factor in fungi downstream of the widely conserved calcium signaling pathway. We find that Crz1 IDRs from anciently diverged fungi can all respond transiently to calcium stress; however, only Crz1 IDRs from the Saccharomyces clade support pulsatility, encode extra information, and rescue fitness, while the Crz1 IDRs from distantly related fungi do none of the three. On the other hand, we find that Crz1 pulsing is conserved in the distantly related fungi, consistent with the evolutionary model of stabilizing selection. Further, we show that a calcineurin docking site in a specific part of the IDRs appears to be sufficient for pulsing and show evidence for a beneficial increase in the relative calcineurin affinity of this docking site. We propose that evolutionary flexibility of functionally divergent IDRs underlies the conservation of stochastic signaling by stabilizing selection.

## Introduction

One of the most remarkable features of living cells is their ability to transmit and process information about their surroundings. It is now appreciated that the dynamics of molecules connected in regulatory networks and signaling pathways underlie many of these capabilities (1– 3). But how do peptide sequences underlying this ability evolve? Relative to enzymatic functions whose evolution has been studied for decades (4–7), research on the evolution of cellular information transmission and signal processing systems is only beginning to emerge (8). Most research on signaling evolution has been focused on the specificity of kinases, receptors, and transcription factors (8), and, to our knowledge, a comparison across species of p53 (9) and Msn2/4 dynamics (Dr. Yihan Lin, personal communication) are the two lone evolutionary studies of stochastic signaling dynamics. Although evolutionary rewiring of DNA-protein interactions in cis-regulatory networks has been described (10), the evolution of the molecular mechanisms that lead to post-translationally controlled signaling dynamics is much less characterized (11–14). Part of the difficulty in obtaining a mechanistic understanding of signaling evolution is that post-translational regulation and transient signaling interactions are often encoded within rapidly evolving IDRs (15–17), which are largely refractory to ancestral protein reconstruction approaches (8,18).

Here we consider the evolution of the pulsatile dynamics of Crz1, a transcription factor in budding yeast that responds to rapid fluctuations of cytosolic calcium concentration (which we refer to as calcium bursts (19)). Pulsatile dynamics are steady state stochastic fluctuations that encode information via frequency modulation of kinase activity (e.g., ERK (20)), protein abundance (e.g., p53(21)), and cytoplasm-to-nuclear-translocalization (e.g., Msn2/4 (22), NFATC1(23,24)). Crz1 has been found to control gene expression through the frequency modulation of post-translationally controlled nuclear-localization pulses (25). To our knowledge, in no case has the fitness benefit of pulsatile dynamics been established, nor has a mechanism for their evolution been proposed, save for one pioneering study of Msn2/4 pulsing (Dr. Yihan Lin, personal communication).

Crz1 is widely conserved in fungi (26,27), but, although the subcellular localization of Crz1 orthologues has been studied in the distantly related fungi *Schizosaccharomyces pombe* (28,29), *Candida albicans* (30,31), and *Cryptococcus neoformans* (32,33), no evidence for pulsing has been reported. Whether Crz1 pulsing is conserved over evolution or whether it is beneficial to the cells has, to our knowledge, not been established. Like other stochastically pulsatile transcription factors, e.g., Msn2 (34) and NFATC1 (35) (Figure 1A), Crz1 contains more than 500 amino acids that are predicted to be intrinsically disordered (Figure 1B, (36)) and contain numerous interaction and post-translational modification sites (e.g., calcineurin docking site, nuclear export signal (NES), nuclear localization signal (NLS), Figure 1B, (37–40)). As expected for an intrinsically disordered region (41), this region shows little primary sequence similarity (Figure 1C).

**Figure 1.**
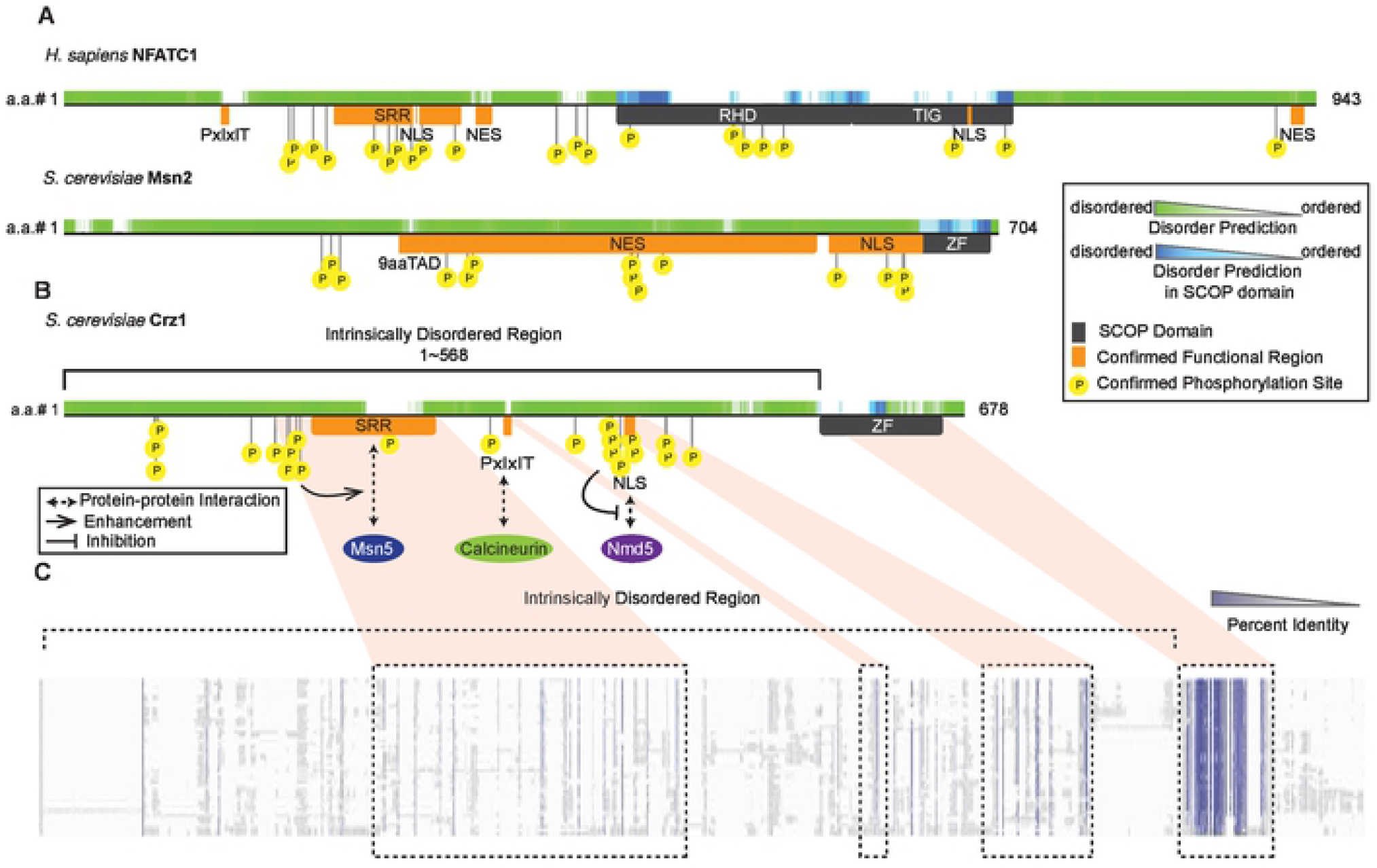
The IDRs of Crz1 and other pulsatile transcription factors contain functional elements involved in the mechanism of nuclear-cytoplasmic translocation. A, B) Schematic representations of three pulsatile transcription factors (NFATC1 and Msn2 in A), Crz1 in B)). The upper half of each representation shows predicted disorder (D2P2, (79)), and the lower half of each representation shows the known functional regions (80,81). SRR: serine-rich region. PxIxIT: *S. cerevisiae* calcineurin docking site PxIxIT. NLS: nuclear localization signal. RHD: Rel homology domain. TIG: transcription factor immunoglobin. 9aaTAD: nine-amino-acid transactivation domain. ZF: zinc finger. B) A schematic drawing of the protein-motif interactions on the protein sequence of Crz1 from *S. cerevisiae*. C) A schematic representation of the sequence alignment of the Saccharomyces fungi from the Yeast Gene Order Browser (YGOB, (82)). Purple shades represent the percent identity of each position (83). Boxes represent the homologous regions of the elements on Crz1 from *S. cerevisiae*. Orange shadows link the corresponding regions between B) and C).

In this study, we show that two orthologous Crz1 IDRs from the Saccharomyces clade support pulsatility, transmit environmental information, and rescue fitness, while two orthologous IDRs from more distantly related fungi do none of the three when expressed in *Saccharomyces cerevisiae*. Furthermore, we show that Crz1 orthologues pulse in the native systems of the two more distantly related fungi, consistent with conservation of the pulsing phenotype over long evolutionary time. The conservation of phenotype but lack of conservation of IDR functions indicates that, within the disordered region, evolutionary changes in some elements needed for complex signaling dynamics have compensated for evolutionary changes in others. This pattern of compensatory evolution in the context of preserved function is a hallmark of stabilizing selection (42). By comparing IDR sequences of Crz1 orthologues, we infer that one of these elements is the calcineurin docking site, PxIxIT, which increased binding strength during evolution. Remarkably, by experimentally increasing the PxIxIT strength in a distantly related IDR to the Saccharomyces PxIxIT strength (via three point-mutations), we can rescue pulsing and improve fitness. Our study demonstrates that stochastic pulsatility is beneficial and that a position-dependent molecular feature in the IDR plays a role in rewiring the molecular basis of the stochastic signaling pathway, even though the phenotype is preserved.

## Results

### Evolutionary changes in Crz1 IDRs are associated with changes in Crz1 pulsing

We first sought to confirm that the IDR of Crz1 was responsible for pulsing. To do so, we designed a passive reporter system in *S. cerevisiae* that expresses an IDR tagged with GFP. We found that the *S. cerevisiae* disordered region alone showed pulsing with similar dynamics as the endogenous protein, although the expression level of the protein was lower (Supplementary Figure 1). Therefore, we fused a defective Crz1 DNA-binding domain (43) tagged with GFP to the disordered regions (denoted as Sc-reporter) and found nearly endogenous dynamics and expression levels, indicating that the disordered region is sufficient for the pulsing dynamics but that the DNA binding domain is needed for protein stability (see methods for more details). Since the calcium/calcineurin signaling pathway is highly conserved (26), we predicted that functional elements within the disordered regions would be conserved over evolution if the dynamics of Crz1 are important for signaling function (25). Consistent with this, some functional elements important for the control of subcellular localization, such as the nuclear localization signal (NLS), nuclear export signal (NES), and the calcineurin docking site PxIxIT (37,38) (summarized in Figure 1B), are found in orthologous Crz1 sequences. On the other hand, overall, the IDRs of Crz1 are highly diverged (little sequence similarity is detected in sequence alignments, Figure 1C), which leads to an opposite prediction that the dynamics of Crz1 orthologues would diverge as has been found for p53 (9). To quantify Crz1 dynamics in response to the upstream calcium signaling pathway, alongside the GFP-tagged Crz1 IDR reporter (denoted as “pulsing reporter”), we expressed a calcium sensor GCaMP3 (44) in a “double-reporter strain” (see methods for more details).

We first confirmed that, as expected based on previous reports for *S. cerevisiae, C. albicans*, and *S. pombe* (28–31), every reporter strain showed transient nuclear localization as a response to 0.2M calcium induction (Figure 2A). However, only Saccharomyces reporters (*S. cerevisiae* (Sc), *Zygosaccharomyces rouxii* (Zr), and *Kluyveromyces lactis* (Kl)) showed pulsing dynamics during steady state (Figure 2B). The two outgroup reporters (*C. albicans* (Ca) and *S. pombe* (Sp)) transiently localized to the nucleus after the calcium induction and then continued with stable nuclear localization during steady state (Figure 2B). To quantify these phenotypic differences at the single-cell level, we measured the duration and amplitude of reporter dynamics by fitting Gaussian Processes to the single-cell trajectories (45). The results suggest that, compared to the outgroup, Saccharomyces reporters have a shorter duration (represented by low ln(l)) and higher amplitude (represented by high ln(a)) in their dynamics (Figure 2C). Because Crz1 pulses are known to follow calcium bursts (19), we adapted the technique of pulse-triggered averaging (46) to investigate the average dynamics of pulsing reporters around calcium bursts. Consistent with the pulsatile dynamics observed in single-cell trajectories, Saccharomyces reporters quickly responded to calcium bursts on average (Figure 2D). In contrast, the average dynamics of the outgroup reporters are not affected by calcium bursts (Figure 2D).

**Figure 2.**
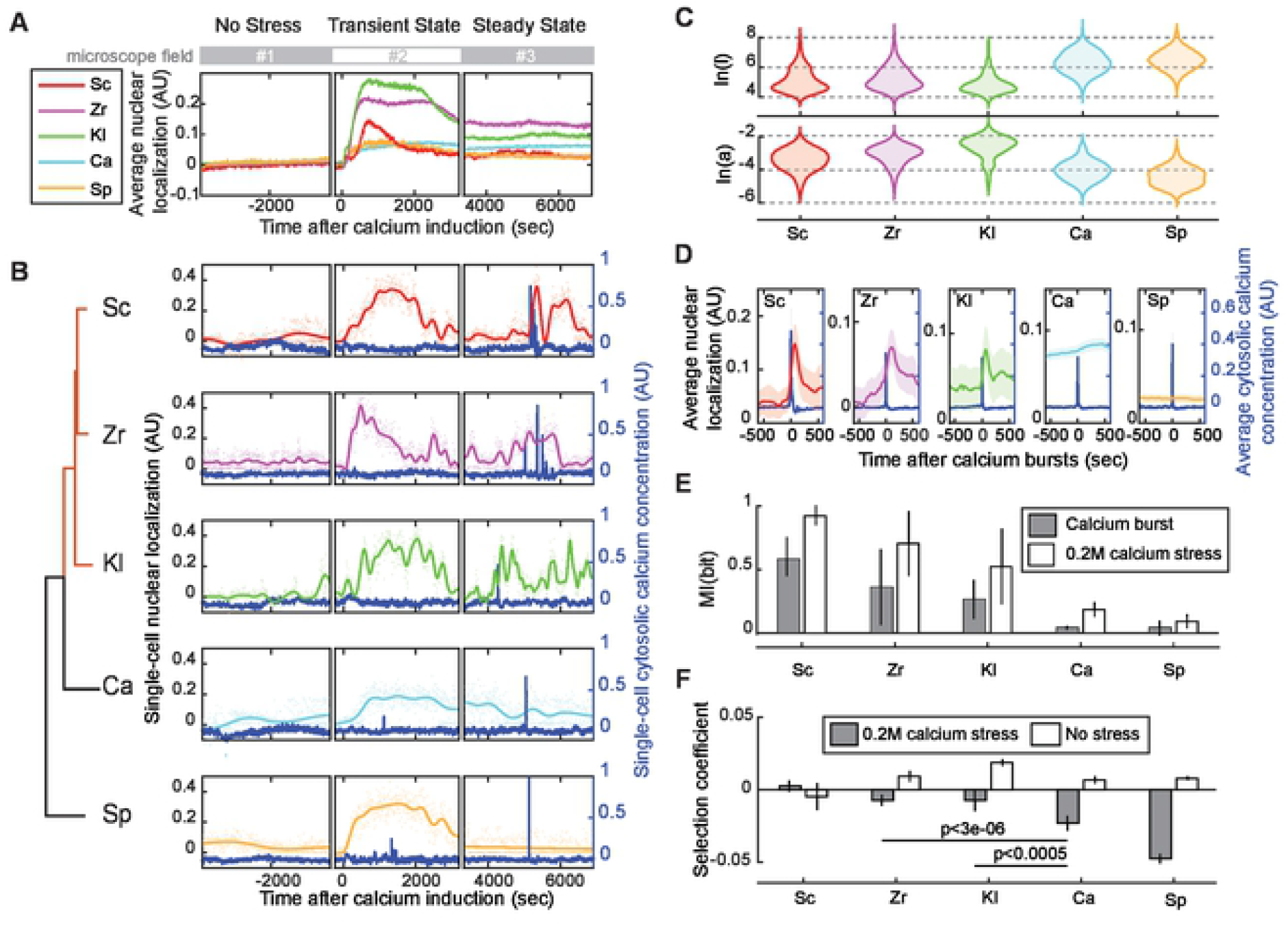
The IDRs of Saccharomyces species rescue both pulsing phenotype and fitness under 0.2 M calcium stress. A) Population averaged localization traces show nuclear localization in response to 0.2 M calcium stress in every reporter strain, corresponding to the Crz1 IDR from the species indicated (Sc, Zr, Kl, Ca, Sp, indicate *S. cerevisiae, Z. rouxii, K. lactis, C. albicans*, and *S. pombe*, respectively). n > 100 cells for each strain. Shadow areas indicate 1.96 SE. Plots are broken to indicate when each experiment switched to a different microscope field to avoid laser-induced nuclear localization. B) Representative single-cell trajectories of cytosolic calcium concentration (blue lines) and nuclear localization (color-coded lines), estimated by Gaussian Process regression on 600 time-points (color-coded dots). Plots are broken to indicate when each trajectory composites a different cell from a different microscope field. The reporter strains (indicated by species names as in A) are ordered according to a phylogenetic tree showing the Saccharomyces clade (red) and the outgroup (black). C) The distribution of dynamic parameters estimated from single-cell time-lapse data. *a* determines the average distance of the trajectory away from its mean, and *l* determines the length of the fluctuation on the trajectory. D) Pulse-triggered averaging of the trajectories of pulsing reporters (black lines) around calcium bursts (blue lines). Time is relative to calcium bursts. Shadow areas indicate 1.96 SE. n > 100 bursts for each strain. E) Averaged mutual information between calcium bursts and dynamics of pulsing reporters (grey bars) and between 0.2M stress and GP parameter values (white bars). n = 1, 3, 3, 3, 1 replicates for Sc, Zr, Kl, Ca, Sp, respectively. F) The selection coefficient measured under 0.2M calcium stress (grey bars) or no stress (white bars). Error bars represent 1.96 SE. P-values indicated are from a two-tail two-sample t-test. n > 10 replicates for each competition assay with at least three cell lines.

Next, we estimated the amount of information encoded in the dynamics of pulsing reporters. Information-theoretic approaches provide a natural framework to estimate the information transmission capacity of cellular signaling pathways (47,48). We estimated the mutual information between the dynamics of cytosolic calcium concentration and pulsing reporters (Figure 2E grey bars). We found that the dynamics of the Saccharomyces reporters encoded more mutual information than that of the outgroup reporters (mean MI = 0.41 bits vs. 0.04 bits, 2-tails t-test, p =0.004, n = 9 and 6, respectively). By analyzing the inferred parameters that describe the steady state dynamics, we also estimated mutual information between the presence or absence of external calcium stress and the dynamics of pulsing reporters (Figure 2E white bars) and again found more mutual information in the Saccharomyces reporters (mean MI = 0.72 bits vs. 0.13 bits, 2-tails t-test, p <10^−3^, n = 9 and 6, respectively). Taken together, these results suggest that pulsing dynamics encode additional information about the environment and that some functional sequence properties arose in the Crz1 IDR along the lineage leading to the Saccharomyces.

### Pulsing confers a fitness benefit in 0.2M calcium stress

If natural selection favored the evolution and preservation of pulsing along the lineage leading to Saccharomyces, pulsing might confer a growth benefit. On the other hand, all the orthologous IDRs support calcium-induced transient nuclear localization (Figure 2A and B, (28,31)) and contain consensus calcineurin docking sites (49), a serine-rich NES region (28–31), and several conserved phosphorylation sites (12) (Figure 1C). Perhaps these are sufficient for cell fitness, and pulsing is simply a non-functional elaboration of this phenotype. Therefore, we sought to directly measure cell fitness under calcium stress (see Methods). To confirm that Crz1 function is needed for fitness in our assay conditions, we tested a mutant with *CRZ1* deletion as well as a mutant that conserved phosphorylation sites in the serine-rich NES region are mutated (“mSRR” (38)). As expected, we found a large fitness defect for the *CRZ1* deletion, and a small but significant fitness defect for mSRR strain, confirming that our assay has the power to detect both large and small effects on fitness (Supplementary Figure 4). The fitness defect of the mSRR strain is also consistent with the functional roles of pulsing (2) because it responds to calcium stress by moving to the nucleus but does not pulse (Supplementary Figure 4). This result indicates that the benefit of pulsing is not simply due to high levels of nuclear localization during steady states.

We next tested whether pulsing is beneficial to the cells by replacing the endogenous IDRs of Crz1 with orthologous sequences. Consistent with the model where pulsing is preserved because it is beneficial to the cell, we found that IDRs from the Saccharomyces clade, but not from the outgroups, rescued fitness in this assay (Figure 2F). The observed fitness defects are conditional on the 0.2M calcium stress (Figure 2F), consistent with the known functions of Crz1(25) and ruling out misfolding or misexpression effects. Although the IDRs of both outgroup species *C. albicans* and *S. pombe* support transient nuclear localization after calcium exposure (Figure 2A), mutants with the outgroup IDRs showed fitness defects comparable to the phosphorylation site mutants (Supplementary Figure 4), indicating that transient nuclear localization is not sufficient for rescuing fitness. Together, these results suggest that pulsing transmits important extra information when cells are under calcium stress that is beneficial for cell growth.

### Pulsing has been under stabilizing selection, but the underlying mechanisms have changed

We next asked if Crz1 pulsing can be found in the native systems of the outgroups. Based on the results above and the lack of previous reports of pulsing in the other species (28–31), we hypothesized that pulsing evolved along the lineage leading to the Saccharomyces from a non-pulsing ancestor. Since we found that pulsing is beneficial to yeast cells under calcium stress, this model is consistent with an adaptive gain of a complex dynamic phenotype. Under this model, we would not expect the outgroup species to show Crz1 pulsing. On the other hand, if selection preserved the pulsing phenotype, we would expect to find pulsing in the other species. Under that model, our finding above that the outgroup IDRs do not pulse in *S. cerevisiae* implies that there must be compensatory changes that maintain pulsing in the outgroup species (42). To distinguish between these models, we obtained strains ((29) and methods) of the two outgroup fungi (*C. albicans* and *S. pombe*) that express endogenous GFP-tagged Crz1 orthologues (CaCrz1 and Prz1) and measured pulsing under 0.2M calcium stress. Remarkably, both CaCrz1 (Figure 3A, B, Supplementary Movie 1) and Prz1 (Figure 3C, D, Supplementary Movie 2) pulse in *C. albicans* and *S. pombe*, respectively, ruling out our hypothesis that pulsing evolved along the lineage leading to the Saccharomyces.

**Figure 3.**
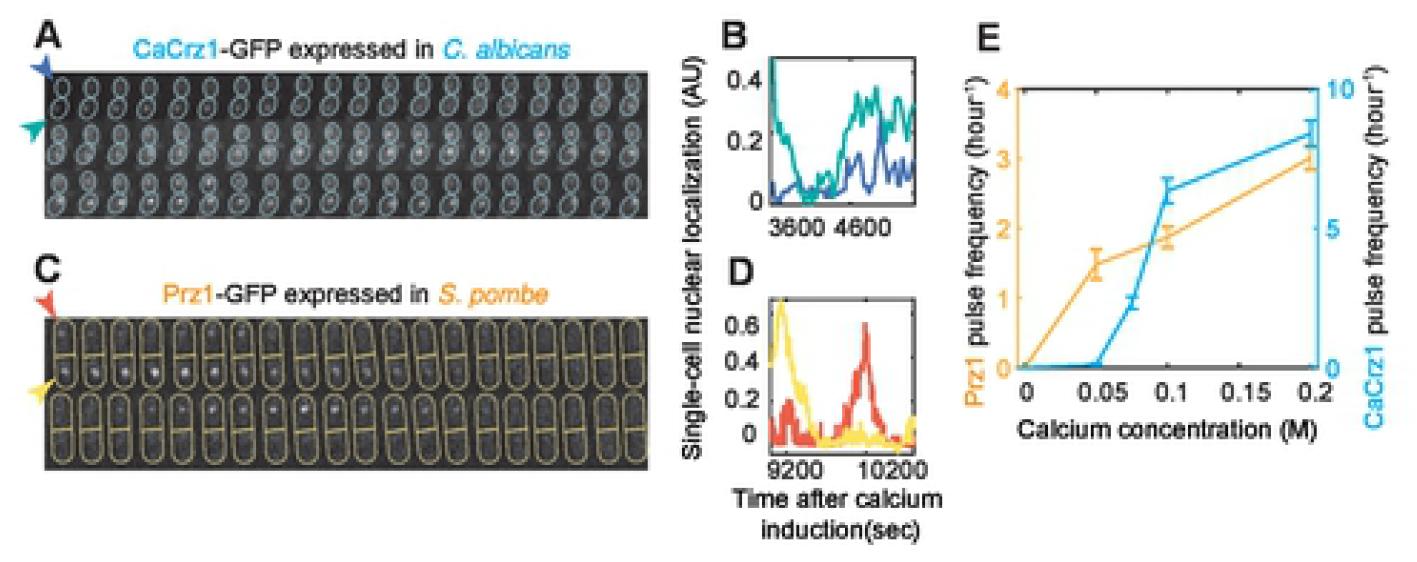
*C. albicans* and *S. pombe* show pulsing and frequency modulation. Filmstrips are showing *C. albicans* (A) and *S. pombe* (C) with GFP-tagged Crz1 orthologues (CaCrz1 and Prz1, respectively) during steady state after the addition of 0.2 M extracellular calcium. Yeast cells are outlined in each frame and indicated through the arrows in the first frame, which are colored accordingly to the single-cell trajectories in B) and D). A) Frames are separated by 30 seconds, and the actual time resolution in B) is the same. C) Frames are separated by 45 seconds, but the actual time resolution in D) is 6 seconds per frame. Image acquisitions of each species were performed through different microscopes. E) Populational average of pulse frequency increases with calcium concentration for both Crz1 orthologues. Error bars represent 1.96 SE. n > 30 in each experiment.

Pulsing of Crz1 in *S. cerevisiae* shows frequency modulation (25), where the rate (per unit time) of pulsing increases with greater calcium stress. Therefore, we measured pulsing in the outgroup species at several calcium concentrations, and, consistent with conservation of frequency modulation, we also found a correlation between CaCrz1 and Prz1 pulse frequency and the strength of calcium stress (Figure 3E). We note that this observation rules out the possibility that our observations of pulsing in these other strains are an artifact of microscopy or laser stress (50). The results are consistent with the model that Crz1 pulsatility (and frequency modulation) has been conserved by natural selection for a long evolutionary time, which is also consistent with our observation of a fitness benefit for pulsing in *S. cerevisiae*. However, the lack of pulsing of outgroup IDRs (and failure to rescue fitness under calcium stress) when expressed in *S. cerevisiae* implies that protein-protein interactions between the IDRs and the calcium signaling pathway has been rewired in some way.

### PxIxIT strength in a specific part of the Crz1 IDR increased in the Saccharomycetacea clade

Because only the IDRs of the Saccharomyces clade support pulsing, we asked which parts of the IDRs are responsible for pulsing. Previous research showed that increasing the affinity of one of the calcineurin docking sites, PxIxIT, leads to a higher pulsing frequency (25). Therefore, we hypothesized that the PxIxIT strength of the Saccharomyces clade is higher than its sister clade that includes *C. albicans* and that this increased strength is needed for pulsing. To test this, we calculated the PxIxIT strength of Crz1 IDRs from 40 fungi of the Saccharomyces clade and the sister clade by using a Position Specific Scoring Matrix (PSSM (51,52), see methods) to predict PxIxIT strength (Supplementary Figure 5A, R^2^ = 0.74 between the measured affinity (K_d_) and the predicted PxIxIT strength of experimentally confirmed PxIxITs (52)). Consistent with the conservation of calcineurin regulation of Crz1, most fungi contain at least one strong PxIxIT site somewhere in their IDRs (Maximum PxIxIT strength >7, Figure 4), indicating the PxIxIT strength alone cannot explain the difference in pulsing between Saccharomyces IDRs and the outgroups.

**Figure 4.**
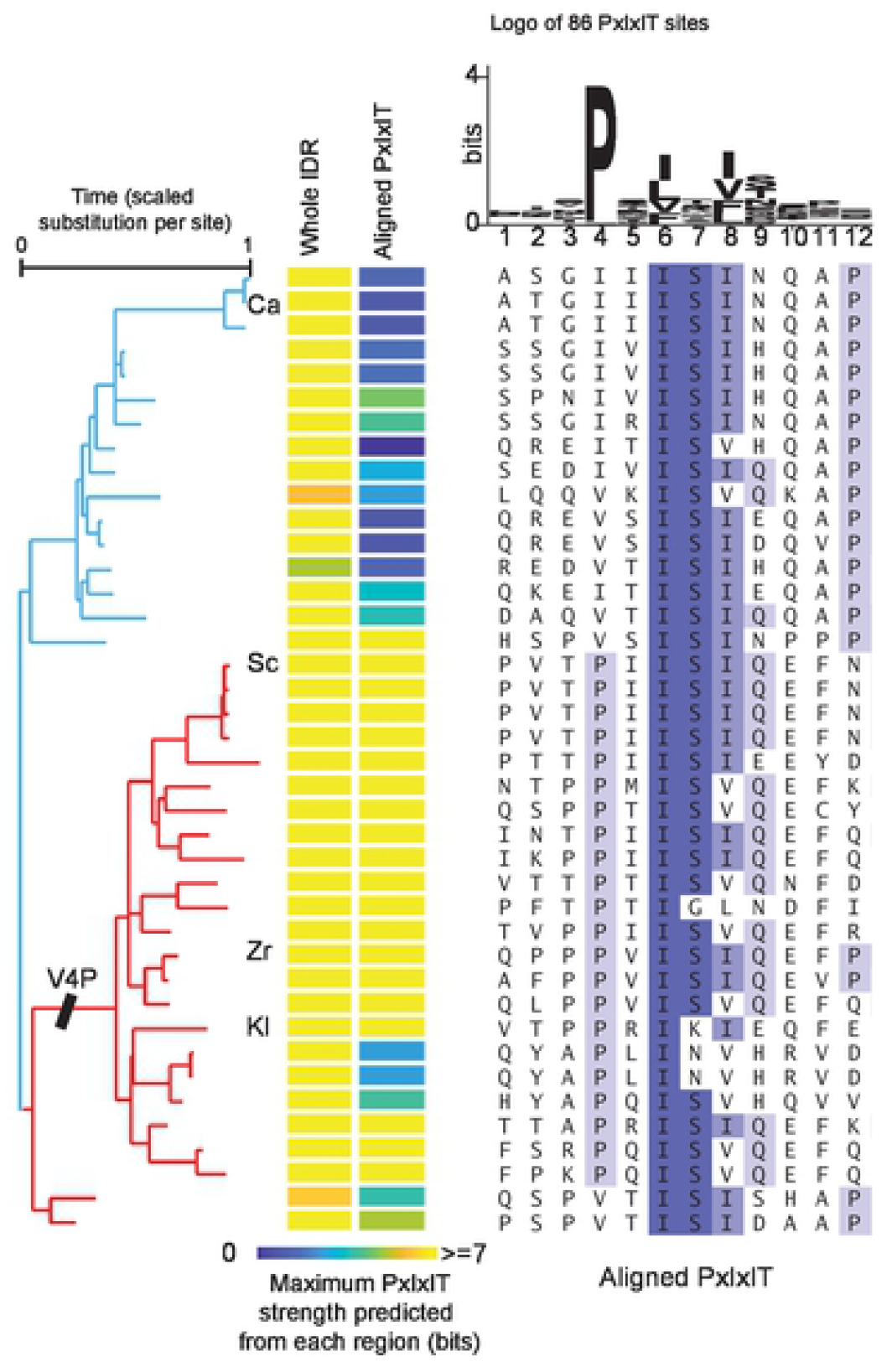
PxIxIT strength in the region homologous to the *S. cerevisiae* docking site is predicted to increase in the lineage leading to the Saccharomyces. The left heatmap represents the strength of the strongest PxIxIT site identified in the whole IDR. The right heatmap shows the predicted strength in the homologous region of the *S. cerevisiae* PxIxIT (shown in the alignment). The branch lengths of the phylogenetic tree are estimated by maximum likelihood based on alignments of the entire Crz1 protein. The label V4P indicates the branch where the inferred V4P substitution occurred. The alignment shows the 12 residues of 40 fungi. The logo represents the PSSM used.

Previous studies showed a connection between the position of calcineurin docking site and dephosphorylation rate (53), suggesting the possibility that evolution can fine-tune some biological functions of largely disordered proteins (e.g., Crz1, NFAT) through selection on the sequence position of a calcineurin docking site. When we aligned the 40 IDRs (54), we found that the *S. cerevisiae* PxIxIT shows a high percent identity in three of the core residues (IS[IV], Figure 4B) but not the proline. The PxIxIT binding pocket of calcineurin buries the proline of the PxIxITs in hydrophobic residues (15,49,53), and as expected, the PSSM shows that proline is highly conserved in confirmed PxIxITs (Figure 4). We inferred a V-to-P substitution on the lineage leading to the Saccharomyces clade, which increases the predicted PxIxIT strength by ∼4 bits (Figure 4), corresponding to a predicted reduction in K_d_ by ∼400μM (Supplementary Figure 5A). Therefore, we hypothesized that the increased strength of the *S. cerevisiae* PxIxIT leads to pulsing and the associated fitness benefit. Because rapidly evolving disordered regions are difficult to align, to rule out the possibility that species outside of the Saccharomyces actually do have a strong homologous PxIxIT site, we repeated this analysis using a 100-residue window around the *S. cerevisiae* docking site and found similar results: in this region of the IDR, only the Saccharomyces showed PxIxIT sites comparable in strength to *S. cerevisiae* (Supplementary Figure 5B)

### Increasing PxIxIT strength in the homologous region of S. cerevisiae IDR is sufficient to rescue pulsing phenotype and fitness

We next tested if the increased PxIxIT strength in the homologous region of *S. cerevisiae* IDR is required for pulsing. We used time-lapse microscopy to investigate if the *S. cerevisiae* PxIxIT can introduce pulsatility into an outgroup IDR. First, we designed a chimeric IDR with the *S. cerevisiae* PxIxIT and C-terminal sequences but the N-terminal region of the *C. albicans* IDR (Figure 5A, denoted as Ca:PxIxIT:Sc). We note that, consistent with the analysis of PxIxIT strength above (Figure 4), the *C. albicans* IDR contains an additional strong PxIxIT in the N-terminal (PSIVIR, a.a.# 23∼28), so this chimera actually contains two strong docking sites. Next, we made a chimera where we swapped the C-terminal portion of the *S. cerevisiae* IDR without the *S. cerevisiae* PxIxIT site. This chimera still retains the N-terminal *C. albicans* PxIxIT site (denoted as Ca:Sc, Figure 5A). Finally, we simply increased the predicted PxIxIT strength in the homologous region to the *S. cerevisiae* level via three point-mutations (Q445T, I446P, N451Q, denoted as Ca^High^, Figure 5A). This construct also contains two strong docking sites.

To determine if chimeric IDRs support pulsatility by responding to calcium bursts, we estimated the mutual information about the presence of calcium bursts from the dynamics of pulsing reporters (Figure 5B, C). We found that the dynamics of both constructs with the *S. cerevisiae* docking site (Ca^High^ and Ca:PxIxIT:Sc) encoded more mutual information than that of the outgroup IDRs (mean MI = 0.21 bits and 0.26 bits, 2-tails t-test, p=0.00577 and 0.00157, n = 3 and 3, respectively). In contrast, we did not find any evidence for the mutual information encoded in the dynamics of Ca:Sc (MI = -0.003 bits, n =1). These results indicate that the *S. cerevisiae* PxIxIT strength is sufficient for the *C. albicans* IDR to support Crz1 pulsatility.

**Figure 5.**
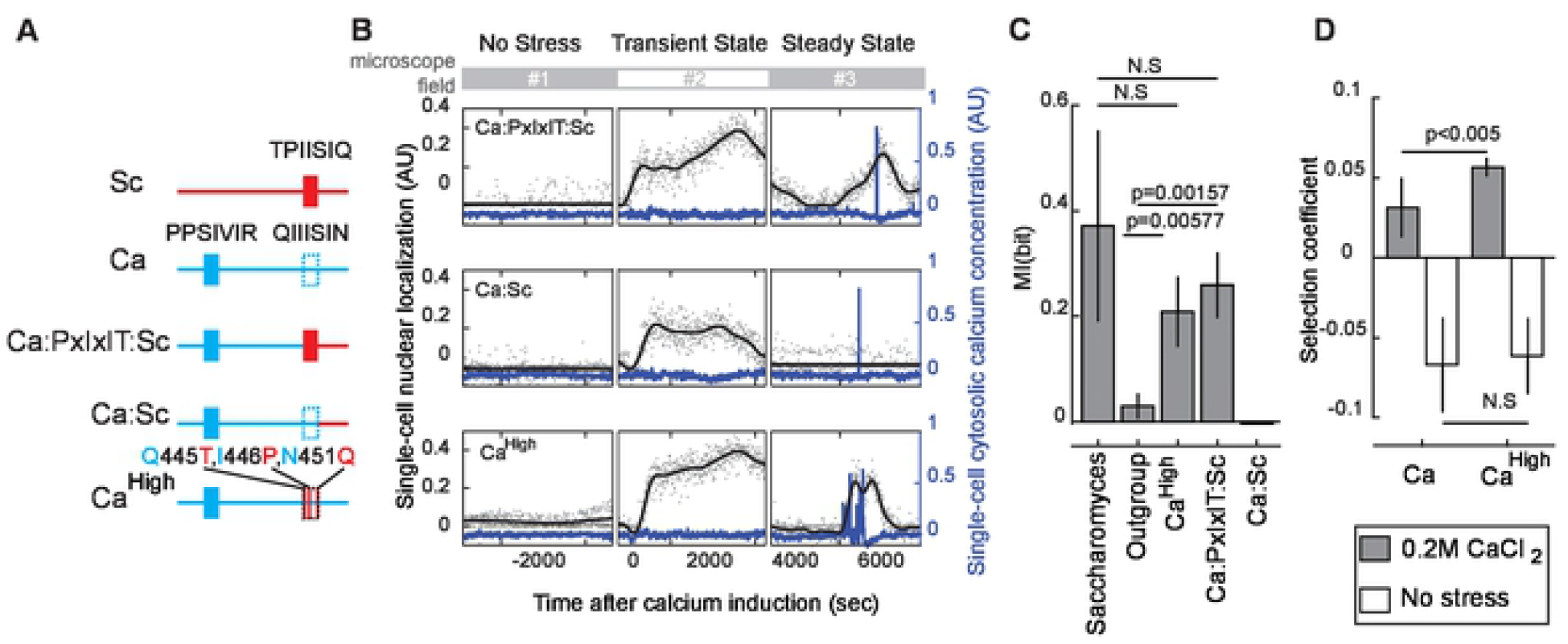
The *S. cerevisiae* PxIxIT strength in the specific part of IDR is sufficient for the *C. albicans* IDR to show a pulsing phenotype and improve fitness under 0.2M calcium stress. A) A schematic diagram of the PxIxIT sites on orthologous and chimeric IDRs. Sc: *S. cerevisiae* IDR; Ca: *C. albicans* IDR; Ca:PxIxIT:Sc: chimeric IDR of the N-terminal region of the *C. albicans* IDR (a.a. #1 to 444) and the C-terminal region of the *S. cerevisiae* IDR including PxIxIT site (a.a. #330 to 568); Ca:Sc: chimeric IDR of the N-terminal region of the *C. albicans* IDR (a.a. #1 to 451) and the C-terminal region of the *S. cerevisiae* IDR excluding PxIxIT site (a.a. #337 to 568); Ca^High^: *C. albicans* IDR with three point-mutations (Q445T, I446P, N451Q). B) Representative single-cell trajectories of cytosolic calcium concentration (blue lines) and nuclear localization (black lines), which are the Gaussian Process regression based on 600 time-points (black dots). Plots are broken to indicate when each trajectory composites a different cell from a different microscope field. C) Averaged mutual information between the calcium burst and the dynamics of pulsing reporters. P-values indicated are from a two-tail two-sample t-test. n = 7, 4, 3, 3, 1 experiments. Error bars represent 1.96 SE. D) The selection coefficient calculated from the competition assays under no stress or 0.2M calcium stress. n = 6 replicates for Ca and 16 replicates for Ca^High^ with three cell lines. Error bars represent 1.96 SE. P-values indicated are from a two-tail two-sample t-test.

Motivated by our previous finding that Saccharomyces IDRs rescue fitness by transmitting environmental information, we wondered whether the extra information encoded in the pulsatile dynamics of the Ca^High^-IDR improves fitness. Therefore, we performed the fitness assay with and without 0.2 M calcium stress and compared the relative fitnesses rescued by Ca^High^-IDR and the *C. albicans* IDR. Previous research showed that Crz1 from *C. albicans* increased the growth rate of the *S. cerevisiae CRZ1* deletion strain (30). Consistent with this, we found that cells expressing the *C. albicans* IDR showed a positive selection coefficient relative to *CRZ1* deletion strains (Figure 5B). Remarkably, the selection coefficient of Ca^High^-IDR expressing cells, relative to the same *CRZ1* deletion strains, was significantly higher than that of the cells expressing the *C. albicans* IDR (mean s = 0.057 vs. 0.031, 2-tails t-test, p=0.0031, n = 16 and 6). In contrast, we did not find a significant difference between the selection coefficients in the absence of stress. These results support our hypothesis that the extra information encoded in the pulsatile dynamics improves fitness. Taken together, our experimental data suggest that the increase in the PxIxIT strength of ancestral IDRs is one way evolution can rewire the signaling pathway while conserving Crz1 pulsing (Figure 6).

**Figure 6.**
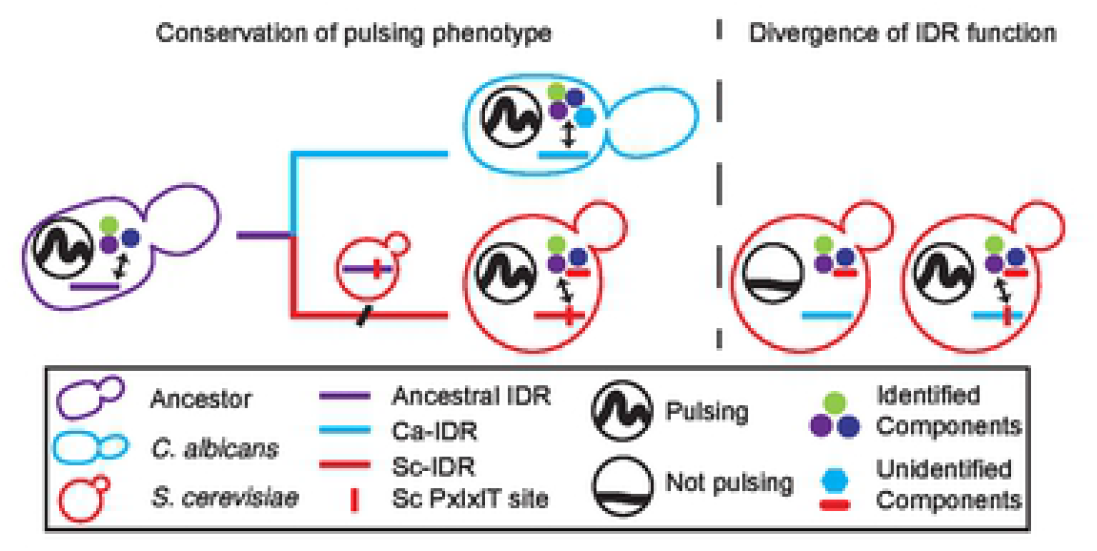
A schematic representation of the evolutionary model that Crz1 pulsing is conserved by stabilizing selection while the underlying mechanism of pulsing has been rewired. Identified components on the signaling pathway (i.e., calcineurin, karyopherin, represented by circles of different colors) are likely to interact with both Ca- and Sc-IDRs, and unidentified components (cyan and red shapes) are likely to interact specifically with each IDR.

## Discussion

Together, our results suggest that Crz1 pulsatility transmits information beneficial for cell growth under 0.2M calcium stress and can be realized by highly divergent and functionally different IDRs. Because of the beneficial effect of transmitting more information through pulsing, our data are consistent with the idea that PxIxIT strength changes were one of the functionally compensatory changes in the calcium/Crz1 signaling pathway under stabilizing selection (Figure 6) (42). Although we cannot rule out convergent evolution that Crz1 pulsing emerged independently in the lineages leading to *S. cerevisiae, C. albicans*, and *S. pombe*, to us, this is an evolutionary model much less parsimonious than that of stabilizing selection. To our knowledge, our results are also the first molecular description of the evolutionary flexibility of post-translationally controlled pulsatility and stochastic dynamic signal processing. Furthermore, our observations illustrate how small evolutionary changes in IDRs can lead to (at least one) functional difference in the mechanism underlying stochastic signaling, ruling out the idea that the rapid sequence divergence is simply due to changes in non-functional residues. More generally, our results are consistent with the idea that IDR sequences encode important functional information and are not “junk proteins” that evolve entirely randomly (55), but they evolve under stabilizing selection (42,56) and accumulate rapid divergence at the sequence level due to the weak constraints relative to folded protein domains (41).

Although the *C. albicans* IDR contains a PxIxIT consensus site in the N-terminal region, pulsing is only supported in *S. cerevisiae* when the PxIxIT strength of the homologous region to the *S. cerevisiae* PxIxIT (a.a.# 445 to 451) is increased, suggesting that calcineurin-dependent pulsatility depends on the position of PxIxIT. Effective dephosphorylation could be necessary for calcineurin-dependent pulsatility(19,23), and previous research has shown instances that the efficiency of dephosphorylation by calcineurin was affected by the distances between calcineurin docking sites and phosphorylated residues (53). Hence, we suggest that evolution has fine-tuned signaling dynamics through the poorly understood position dependency of short linear motifs in this case.

Previous quantitative studies on dynamic signal processing focused on the information encoded by synchronous, transient dynamics (47,48,57). Hence, it remained unclear whether stochastic pulsatile dynamics during steady state transmits information that is beneficial for cell growth. Because all five IDRs in our study show transient nuclear localization after calcium induction (Figure 2A), we could measure the effect of steady state stochastic pulsing while minimizing the effects of differences in the transient dynamics. We not only show that the stochastic dynamics during steady state encode environmental information but also that information is transmitted between two stochastic components on the same pathway: the bursting dynamics of cytosolic calcium concentration and the pulsatile dynamics of Crz1 nuclear localization. To our knowledge, this is the first application of information theory to show information transmission between two unsynchronous and stochastic cellular dynamics.

## Methods

### Yeast strains

Lasers used during fluorescence microscopy are known to induce nuclear localization of Crz1 and can affect the dynamics of the pulsing reporters (50). To minimize these effects but still measure dynamics of both cytosolic calcium concentration and Crz1 nuclear localization, we designed double-reporter strains expressing the pulsing reporters described in the text (IDRs followed by yEGFP-tagged defective zinc finger, IDR-dZF-yEGFP) and the calcium reporter GCaMP3 (44). Plasmids expressing the pulsing reporters were constructed using Gibson assembly protocol (58). The pulsing reporter genes were assembled between the promoter of CRZ1 and the ADH1 terminator (pCRZ1-IDR-dZF-yEGFP-tADH1) and integrated at the *URA3* locus of reference strain BY4741 using a selectable marker (URA3). The fragments of orthologous IDRs were amplified from the genomic DNA of the corresponding species and corrected the CUG codon usage (59). The IDR with the mSRR mutations was constructed by modifying the Sc-pulsing reporter plasmid (URA3::pCRZ1-ScIDR-dZF-yEGFP-URA3MX). The chimeric IDRs were constructed through two-fragment transformations, with each fragment amplified from the Sc-pulsing reporter plasmid (URA3::pCRZ1-ScIDR-dZF-yEGFP-URA3MX) or Ca-pulsing reporter plasmid (URA3::pCRZ1-CaIDR-dZF-yEGFP-URA3MX). In the same strains, we integrated a previously constructed pRPL39-GCaMP3-tADH1 at the *HO* locus using a selectable marker (LEU2) (19). All transformations were performed using the standard lithium acetate procedure (60).

Compared to the previously constructed dual-color strains (19), the *S. cerevisiae* double-reporter strain required ∼90% lower laser intensity (no observable nuclear localization induced by laser stress (50)) to record both the dynamics of cytosolic calcium concentration and Crz1 nuclear localization. Because of the spatial differences in the patterns (nuclear Crz1-GFP vs. cytoplasmic GCaMP3), the two signals can be distinguished using a two-component mixture model (described in the methods section of Reporter intensity quantification).

Each fitness assay strain was constructed by integrating an IDR and a wild type zinc finger at the endogenous locus of CRZ1 through two-fragment transformation using a selectable marker (HIS3). The zinc finger was tagged with yEGFP to report expression level. The IDR fragments were amplified from the existing plasmids expressing the corresponding pulsing reporters and zinc fingers or amplified from the genomic DNA of the corresponding double-reporter reporter strains. For the competition assay on plates (see below), each fitness assay strain was labeled with red by genomic integration of pRPL39-yemCherry-tADH1 at the promoter region of CAN1 using a selectable marker (LEU2).

The GFP-expressing *S. pombe* strain is a gift from Dr. Gordon Chua (29).

The *C. albicans* strain CaLC7415 with both copies of *CRZ1* C-terminally tagged with GFP was made using a transient CRISPR approach adapted from Min *et al*. (61). The GFP-NAT cassette was PCR amplified from pLC389 using oLC9367 and oLC9368 (see tables below for plasmids and oligos). The CaCAS9 cassette was amplified from pLC963 using oLC6924 and oLC6925. The sgRNA fusion cassette was PCR amplified from pLC963 with oLC5978 and oLC9371 (fragment A) and oLC5980 and oLC9372 (fragment B), and fusion PCR was performed on fragments A and B using the nested primers oLC5979 and oLC5981. The GFP-NAT cassette, sgRNA, and Cas9 DNA were transformed into SN95. Upstream integration was PCR tested using oLC600 and oLC9369, and downstream integration was tested using oLC274 and oLC9370. Lack of a wild-type allele was PCR tested using oLC9369 and oLC9373.

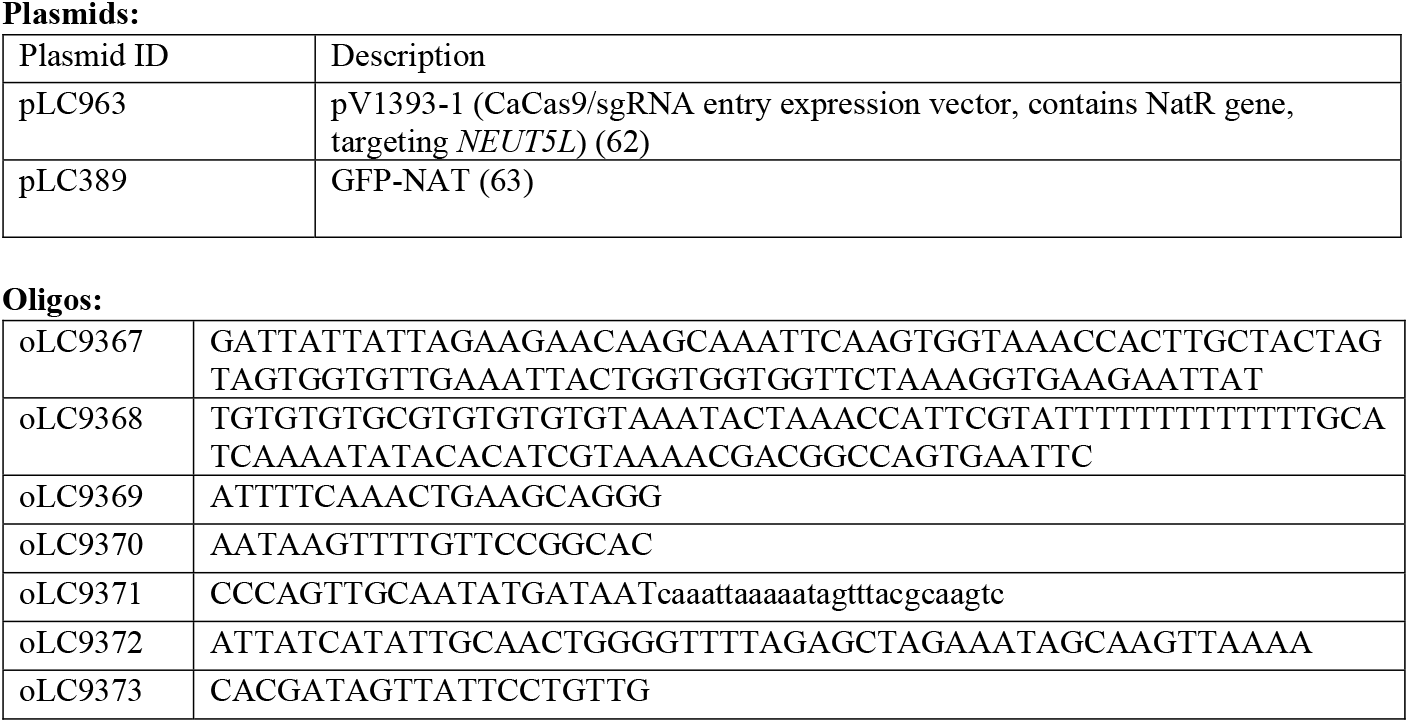

### Spinning-disk confocal microscopy and image analysis

A Nikon CSU-X1 was utilized for time-lapse imaging at room temperature (22n) for *S. cerevisiae* and *S. pombe* strains. 488 nm laser was applied with time resolutions of 6 sec/frame, exposure time of 50 msec, and 25% laser intensity. Bright-field images with out-of-focus black cell edge were acquired every minute for cell segmentation and tracking. The growth conditions were based on a standard protocol (19,29). A Zeiss Axio Observer was utilized for time-lapse imaging at room temperature (22□) for *C. albicans* strain. 488 nm laser was applied with time resolutions of 30 sec/frame, exposure time of 100 msec, and 100% laser intensity. All the time-lapse imaging experiments were started when cells were in log-phase. Cells were cultured in YPD with a carbon source of 2% glucose overnight and in SC during time-lapse imaging. Time-lapse movies of every strain had been replicated on different days to control day-to-day variation and showed reproducible results.

Segmentation was automatically performed by YeastSpotter (64). Cell tracking was performed by identifying 90% of overlapping cell areas between two frames. Mis-segmented and miss-tracked objects were manually removed. 100-300 cells were identified in each time-lapse movie. Single-cell photobleaching correction was conducted after single-cell reporter intensities were quantified (see below) using bi-exponential regression (65) with the baseline of the calcium reporter.

### Reporter intensity quantification

For each time point, The cytosolic intensity of the calcium reporter and the nuclear intensity of the pulsing reporter was quantified by fitting a mixture of a Gaussian distribution and a uniform distribution to the pixel intensities of each segmented cell area, and the parameters of distributions were estimated using expectation-maximization (see the supplementary text of (19) for more details and derivation of the algorithm). Once the parameters were estimated, the estimate of the calcium reporter at a time point is the mean of the Gaussian distribution, and the estimate of nuclear localization is the difference between the means of the uniform distribution and the Gaussian distribution. The algorithm can reproduce previous observations from time-lapse movies where Crz1-RFP and GCaMP3 are merged into one channel (Supplementary Figure 2).

### Competition fitness assay on plates

Previous research showed that competition assays could be robustly performed through plate readers (66), which provides both the sensitivity of competition assays (67,68) and the high performance of growth assay on plates (69). We followed this approach and recorded competition of fitness strains with a plate reader. Cells were grown in SC media at 30□ for ∼48 hours and then serially diluted to 1/1024 of the initial concentration on a flat-bottom 96-well plate. The plate reader Tecan M1000 was automatically run by the application Tecan i-Control. OD600 and RFP intensity were measured every 15 minutes for 24 hours at 30□, and the plates were constantly shaken through the whole experiment. Similar to the protocol of competition-based fitness assay on plates (66), the wells on the plates were either monoculture or mixed-culture. The mono-cultural wells contained only the RFP-labeled strain and were for the calibration between RFP intensity and OD600 absorbance through linear regression. The mixed-cultural wells contained both the RFP-labeled strain and the colorless strain. The absorbance of the colorless strains in the mixed-culture wells was estimated by subtracting the measured absorbance by the absorbance inferred from the RFP intensity, so the growth curves of both competing strains in each mixed-culture well can be obtained. The time point when the growth curves reached the 10^th^ generation was identified and kept consistent throughout each experiment. The relative selection coefficients were calculated with the formula (70,71)

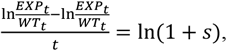

where *t* means the number of generations and *s* is the selection coefficient.

We found that this approach provides a resolution of the selection coefficient to 10^−3^ and successfully reproduced a previously reported small fitness defect (Supplementary Figure 3).

### Single-cell trajectory quantification with Gaussian Process

In the previous studies of pulsatile transcription factors, pulses were identified before quantification and statistical analysis, e.g., pulse frequency(20,25) and pulse triggered averaging (46). This approach presumes that the dynamics are pulsatile. However, in our case, whether a pulsing reporter pulse or not was to be determined. Therefore, we needed a more general approach to quantify single-cell trajectories.

We used a Gaussian Process regression model (19,45,72) with the squared exponential kernel to summarize each single-cell trajectory. The kernel can be expressed as

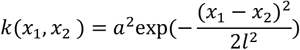

where *x*_1_, *x*_2_ indicate a pair of nuclear localization scores at different time points, *a* determines the average distance of the trajectory away from its mean, and *l* determines the length of the fluctuation on the trajectory. We used was the default MATLAB (Mathworks) function for the Gaussian process, `fitrgp`. Estimation was considered numerically unstable if ln(*a*) < −6, and cell trajectories were removed if their estimates were below this value.

### Estimation of mutual information and pulse-triggered averaging

Information theory provides a natural framework(73) to quantify information transmission in cells as mutual information (MI). Previous studies estimated MI encoded in signaling pathways (47,48) with decoding methodology (e.g., an SVM classifier (48)). Here we adopted a widely used kNN estimator (k=4, (47,74)) for the advantage of its simplicity. Two different parametric forms of trajectories (described below) were applied to estimate the MI between the calcium stress and the dynamics of pulsing reporters and the MI between the calcium bursts and the dynamics of pulsing reporters.

To estimate MI between the calcium stress and the dynamics of pulsing reporters, single-cell trajectories of pulsing reporters from one experimental replicate were parameterized with the Gaussian process regression model described above. Gaussian Processes normalizes the trajectories to their mean amplitudes, excluding information about the absolute level of nuclear localization. For each environmental condition (no stress or 0.2 M calcium stress), an equal number of trajectories were randomly selected and parameterized. The selected data were processed into *D* = {(**x**_1_, *y*_1_), (***x***_2_, *y*_2_),…, (**x**_*n*_, *y*_*n*_)} that consist of *i* = 1, …, *n* pairs of parameter values, **x**_*i*_ = {*a*_*i*_, *l*_*i*_}, and their corresponding environmental labels of 2 discrete values, y_*i*_ ∈ {*c*_0_, *c*_1_} (no stress or 0.2 M calcium stress). **x** was jittered to avoid identical samples. The estimated MI(**x**; *y*) o one experimental replicate was bootstrapping 60 times for the average value.

To estimate MI between calcium bursts and the dynamics of pulsing reporters, we first categorize time points on each calcium trajectory into two groups. A typical trajectory of calcium reporter contains two types of fluctuations: a baseline of slow fluctuation and calcium bursts as rapid and large fluctuation. Our previous research showed that only calcium bursts lead to Crz1 pulses (19); hence, the information about the presence of calcium bursts should be encoded in a short period of the pulsing dynamics after the bursts. Precisely, let a trajectory of pulsing reporter ***x*** = {*x*_1_, *x*_2_, …, *x*_*p*_} consist of *t* = 1, …, *p* nuclear localization scores, a block of the trajectory *X* = {*x*_*t* + 1_, *x*_*t* + 2_,*…, x*_*t* + τ_} should encode the information about the states of preceding calcium trajectory at time *t* (calcium burst or basal fluctuation) with the mean score ⟨*X*⟩ and the mean velocity of the score 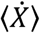. Our goal is to process data extract from j = 1, …, *m* blocks (*X* = {*x*_*t* + 1 +_ τ_(*j* -1)_, *x*_*t* + 2 + τ(*j*-1)_,*…, x*_*t+τj*_}) such that *D*_*j*_ = {(**x**_1*j*_, *y*′_1_),(**x**_2*j*_, *y*′_2_), …, (**x**_*nj*_, *y*′_*n*_} consist of *i* = 1, …, *n* pairs of statistic summaries, 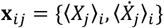, and their corresponding burst labels of 2 discrete values, *y*′_*i*_ ∈ {*c*′_0_, *c*′_1_} (no calcium burst or calcium burst). We then estimate MI_*j*_ (**x**_j_, *y*′) as the information encoded in the *j*th block of pulsing trajectory after a calcium fluctuation. The details of the pipeline are provided in Appendix. We found that the estimation from the first block, MI_1_, is representative of each experimental replicate and reported MI_1_ in the results section.

We use a similar pipeline to adapt pulse-triggered averaging by simply averaging all the trajectories of pulsing reporters in a 20-min window centered around every labeled calcium burst.

### Sequence analyses

To predict the calcineurin docking strength of a sequence, we used a Position Specific Scoring Matrix (PSSM). PSSM is a standard statistical model widely used for predicting transcription factor binding strength of a DNA motif (75) or protein binding strength of a short linear motif in an IDR (51,52,76). In this study, a PSSM was constructed with 86 experimentally confirmed calcineurin binding sites (so-called PxIxIT sites) collected from the ELM database (77) and other sources (26,43,52,78). The calcineurin docking site of Crz1 was excluded to avoid circularity. The docking strength *S* of a sequence *X* with sequence length *w* was calculated as

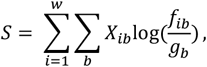

Where *b* **∈ A**. **A** indicates the 20 amino acids, *X*_*ib*_ = 1 if the sequence is amino acid *b* at position and 0 otherwise, *f*_*ib*_ is the probability of observing amino acid *b* at position in a calcineurin docking site (from the PSSM), and *g*_*b*_ is the probability of observing amino acid *b* in the genomic background distribution and was assumed to 1/20. To test how well this simple model predicts measured calcineurin binding affinity, we compared the predicted strength, *S*, to the affinity of 10 characterized PxIxIT sites (52) and found that a linear model where a change of 1 unit of *S* (which is measured in bits) corresponds to 74.4 unit of K_d_ (measured in μM, SE = 15.4, p = 0.001)

## Acknowledgments

Shadi Zabad contributed several analyses that were ultimately not included in the manuscript. We thank Drs P. Beltrao, G. Liti, M. Cyert, and Y. Lin and members of the Moses lab for discussions and helpful suggestions. We thank Drs M. Cyert, Y. Lin, and M. Elowitz for their comments on the manuscript. We thank Drs L. Cowen and G. Brown for the genomic DNA of *C. albicans* and *S. pombe*. We thank Dr. Chua for the GFP-expressing *S. pombe* strains. LEC is supported by the Canadian Institutes of Health Research Foundation Grant (FDN-154288). LEC is a Canada Research Chair (Tier 1) in Microbial Genomics & Infectious Disease and co-Director of the CIFAR Fungal Kingdom: Threats & Opportunities program. This research was supported by an NSERC discovery grant to AMM, a QEII-GSST scholarship to ISH, and infrastructure obtained with grants from the Canada Foundation for Innovation to AMM.

## Figure legends

Supplementary Figure 1. The IDR of Crz1 is sufficient for pulsing. A) Representative images of GFP channel and bright-field channel of three strains expressing endogenous Crz1 tagged with GFP (Crz1-GFP), a passive reporter of the IDR tagged with GFP (zinc fingered removed, ΔZF-reporter), and a passive reporter of *S. cerevisiae* Crz1 with defective zinc fingers (Sc-reporter), respectively. B) Representative trajectories of each strain. The trajectories of nuclear localization (black lines) are the Gaussian process regression based on nuclear localization score of 600 time-points (cyan dots).

Supplementary Figure 2. Two dynamics recorded with two-color images and merged into one-color images can be distinguished via a mixture model. A) An example of single-cell trajectories before (upper plot) and after (lower plot) merging for Crz1 (dots) and calcium (blue trace). Black traces are Crz1 trajectories smoothed with Sacitzky-Golay filtering. Black circles indicate Crz1 pulses. B) Calcium bursts (left plot) and Crz1 pulses (right plot) identified in the experiments sorted from large to small. C) The distributions of the change in the number of identified Crz1 pulses after merging. D) The probability of first, second, third, and fourth Crz1 pulses plotted as a function of the time they occur relative to calcium bursts from the same cells. The left and the right stacked histograms are data from the separate images and the merged images, respectively. E)Data are divided into three groups based on calcium burst sizes and aligned to each group’s mean calcium burst size. The dots’ size represents the probability of finding a number of Crz1 pulses in a group (summed up to 1 in each column).

Supplementary Figure 3. The competition fitness assay on 96-well plates can reproduce the significant growth defect of a mutant (noted as ‘5A’) reported by Zarin et al., 2017. A) Fluorescent intensity of the monoculture. Each marker represents the fluorescent intensity and OD of a well at each time-point. B) Mean selection coefficient calculated from the fluorescent data of mixed culture. Error bars represent 1.96 SE. Dashed line indicates the selection coefficient of 5A mutant reported by Zarin et al., 2017 (−0.038) C) The growth curves of WT and 5A strains in the mixed cultures predicted by the algorithm of Ram et al., 2019. The selection coefficient is -0.05, which is calculated with the first and the last time point of the predicted growth curves.

Supplementary Figure 4. Mutations in the conserved phosphorylation sites in the serine-rich region lead to constant nuclear localization after 0.2M calcium induction and fitness defect under 0.2M calcium stress. A) Upper panels show the population average of nuclear localization score. Shadow indicates SD with n > 100 for each strain. Lower panels show representative single-cell trajectories of cytosolic calcium concentration (blue lines) and nuclear localization (black lines), which are the Gaussian Process regression based on 600 time-points (red dots). Plots are broken to indicate when each experiment switched to a different microscope field to avoid laser-induced nuclear localization. B) The selection coefficient obtained from the competition assays under 0.2M calcium stress (grey bars) or no stress (white bars). Error bars represent 1.96 SE. n > 10 replicates for each competition assay with at least three cell lines.

Supplementary Figure 5. Calculate PxIxIT strength with PSSM. A) Predicted PxIxIT strength plotted against the measured PxIxIT affinity of the same sequences from the database of ref (52). Linear regression model: y ∼ 779-74.4x, R^2^ = 0.74. B) Heatmaps represent the maximum PxIxIT strength calculated from the sub-sequences of the whole IDR or the 100-residue homologous region around the *S. cerevisiae* PxIxIT.

## Appendix

### Titles of supplementery movies

Supplementary Movie 1. An example of CaCrz1 pulsing in *C. albicans*

Supplementary Movie 1. An example of Prz1 pulsing in *S. pombe*

### Details of MI estimation for one experimental replicate

1. For each single cell
  a. Labeling calcium burst:
    i. Fit the trajectory of calcium reporter with the Gaussian Process described above to estimate basal fluctuations
    ii. From the same trajectory, identify the highest 100 peaks (local maxima) with a minimal interval of 30 sec, and classify peaks into calcium bursts or basal fluctuations with the 95% CI of baseline estimated via the Gaussian Process, **Y** = {*y*′_1_, *y*′_2_,…, *y*′_*1oo*_}^*T*^, *y*_*i*_ ∈ {*c* _0_, *c*_1_}. Record the time points of each peak, *T* = {*t*_1_, *t*_2_,…, *t*_100_}, *t*_*i*_ ∈ {1,2, …, *p*}. We used MATLAB function findpeaks to identify peaks and time points.
  b. Process pulsing dynamics of the *j*th block:
    i. Fit the trajectory of pulsing reporters with the Gaussian Process described above to smooth the trajectory
    ii. For every calcium peak *′*_*i*_ the *j*th block was chosen 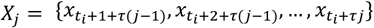. We arbitrarily chose *r = 5*. Calculate 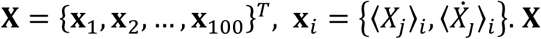 was jittered to avoid identical samples.
2. Concatenate every **X** and **Y** of the cell population. Randomly withdraw an equal number of samples for each burst label and reorganize the data into = {(**x**_1_, *y*′_1_), (**x**_2_, *y*′_2_), …, (**x**_*ni*_, *y′n}*. Estimate MI by bootstrapping 600 times and report mean MI (**x**_*j*_, *y*′).

### List of 40 fungi used for sequence analysis

*Candida dubliniensis*

*Candida albicans*

*Candida tropicalis*

*Candida parapsilosis*

*Candida orthopsilosis*

*Lodderomyces elongisporus*

*Spathaspora passalidarum*

*Millerozyma farinose*

*Scheffersomyces stipites*

*Meyerozyma guilliermondii*

*Debaryomyces hansenii*

*Debaryomyces fabryi*

*Clavispora lusitaniae*

*Candida auris*

*Metschnikowia bicuspidata*

*Babjeviella inositovora*

*Saccharomyces cerevisiae*

*Saccharomyces mikatae*

*Saccharomyces kudriavzevii*

*Saccharomyces uvarum*

*Candida glabrata*

*Kazachstania Africana*

*Kazachstania naganishii*

*Naumovozyma castellii*

*Naumovozyma dairenensis*

*Vanderwaltozyma polyspora*

*Tetrapisispora phaffii*

*Tetrapisispora blattae*

*Zygosaccharomyces rouxii*

*Zygosaccharomyces parabialii*

*Torulaspora delbrueckii*

*Kluyveromyces lactis*

*Eremothecium gossypii*

*Achbya aceri*

*Eremothecium cymbalariae*

*Lachancea kluyveri*

*Lachancea thermotolerans*

*Lachancea waltii*

*Cyberlindnera fabianii*

*Wickerhamomyce ciferrii*

